# Change in Monarch Butterfly Winter Abundance Over the Past Decade: A Red List Perspective

**DOI:** 10.1101/2023.01.12.523862

**Authors:** Timothy D. Meehan, Michael S. Crossley

## Abstract

1. Assessing invertebrate species for the IUCN Red List under Criterion A requires fitting an appropriate statistical model to available abundance data and calculating a ten-year change (TYC) estimate from predicted abundances. When the rate of change has not been constant across the entire time series, models that accommodate variable change rates are strongly recommended.

2. The monarch butterfly (*Danaus plexippus*) was recently added to the IUCN Red List (A2ab Endangered) based on analysis of data on winter abundances in Mexico and the western USA between 1993 and 2020. TYC estimates in the monarch assessment came from models that assumed constant change rates. We conducted a Bayesian analysis of the same data using models that accommodated variable change rates and used those models to compute TYC estimates.

3. Our results suggested that monarch population change rates have not been constant. The analysis yielded a model averaged TYC estimate of +5.23%, which was not statistically distinguishable from 0% and was considerably different from values of −46% and −72% in the assessment. The Bayesian posterior probability of a TYC value below −30% (A2ab Vulnerable) was 0.15 and that of a TYC value below −50% (A2ab Endangered) was 0.03.

4. We suggest that a more thorough analysis of recent overwintering abundances will lead to an improved IUCN assessment for monarch butterflies. We recommend that other researchers evaluating monarch conservation status consider using models with variable change rates, as models with constant change rates may not accurately predict the trajectory of monarch abundances into the future.

## Introduction

Evaluating a species under Criterion A of the IUCN Red List requires an estimate of population change over the past ten years, or three generations, whichever is longer (IUCN, 2022). For invertebrates with short generation times, Red List Guidelines recommend fitting an appropriate statistical model to available census data and calculating a ten-year change (TYC) estimate based on model-predicted abundances as 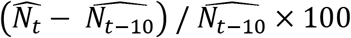, with 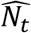 being the predicted abundance for the most recent year (Akçakaya *et al*., 2021; IUCN, 2022). When the available time series is longer than ten years, Red List Guidelines recommend fitting an appropriate model to the full time series and, again, calculating TYC based on model predictions from 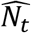 and 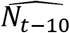 (Akçakaya *et al*., 2021; IUCN, 2022). Given the reliance of TYC estimates on model predictions, it is very important to select a model that properly fits the full time series (Akçakaya *et al*., 2021).

Many types of models can be fit to an abundance time series. If the proportional rate of change has been constant, then a simple exponential change model may suffice (Akçakaya *et al*., 2021; IUCN, 2022). However, when the rate of change has not been constant, more complex models are preferred, including those with logistic functional forms (Akçakaya *et al*., 2021), segmented regressions (Thogmartin *et al*., 2020; Akçakaya *et al*., 2021), or smooth models such as generalized-additive models (Fewster *et al*., 2000; Smith & Edwards 2021) or state-space models (Semmens *et al*., 2016; Auger-Méthé *et al*., 2021). Well established model-fit and – comparison techniques (Hooten & Hobbs, 2015) are recommended to determine the best, or best set of, candidate models from which to derive TYC estimates (Akçakaya *et al*., 2021).

In December 2021, the migratory monarch butterfly (*Danaus plexippus*) was listed by the IUCN as an Endangered Species under Category A2ab, which requires reasonable evidence for a TYC of −50% (IUCN, 2022). The IUCN assessment for the species (Walker *et al*., 2022) emphasized three TYC values: −22%, −46%, and −72%. The value of −22% was specific to monarchs that bred in eastern North America and wintered in Mexico and was derived from a growth rate estimate from Voorhies *et al*. (2019), who described it as the average annual change in winter abundances between 1993 and 2019. It is not clear why a long-term average change rate was considered in the species assessment instead of the change rate calculated from model predictions from the past ten years; the number was presumably derived from a state-space model (Semmens *et al*., 2016) that produces annual population estimates that could have been used to calculate TYC. Uncertainty about that point estimate was not discussed in the assessment, despite large uncertainties being reported in Semmens *et al*. (2016) and Voorhies *et al*. (2019).

The other two TYC values given, −46% and −72%, came from an analysis detailed in the supplemental materials of the species assessment (Walker *et al*., 2022). That analysis used a composite time series of total annual monarch abundance from the wintering region in Mexico (1993—2020) and a wintering region along the west coast of the USA (1997—2020). TYC values of −46% and −72% were derived from simple exponential and linear change models, respectively, fit to the full time series (Figure 1). Quick inspection of the time series suggested that other models might fit the data at least as well as simple exponential or linear change models. Indeed, work published before completion of the assessment (Thogmartin *et al*., 2020) demonstrated that a segmented model with different change rates before and after 2013 or 2014 was superior (by 6 AIC units) to a model with a single, constant change rate. The rate before the change point was clearly negative and the rate after the change point was not different from zero (Thogmartin *et al*., 2020), and this pattern has become more apparent as more data have been added to the time series (Figure 1). As discussed by Akçakaya *et al*. (2021), if growth rates have been variable over a long time series, then predictions from constant change rate models should not be used to compute TYC values. More complex models should be applied to the data, their model fits compared to those of simpler models, and predictions from the best-fitting model(s) should be used to compute TYC estimates for Red List assessment (Akçakaya *et al*., 2021).

**Figure 1.**
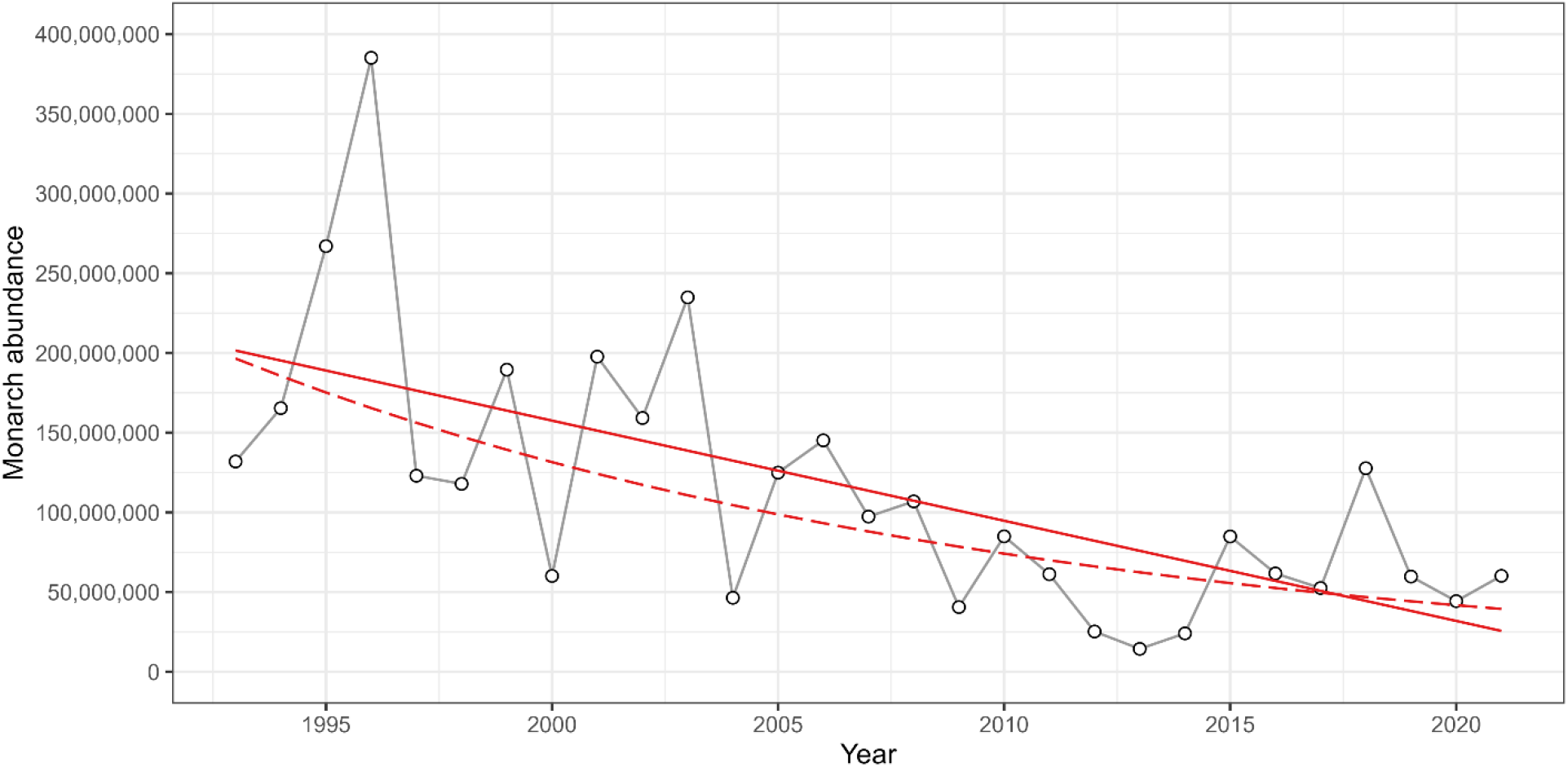
Estimated total winter abundance of the migratory monarch butterfly from 1993 through 2021. The solid red line is the fit of a linear change model. The dashed red line is the fit of an exponential change model. Note the pattern where red lines tend to over predict from 2007 through 2014 and under predict from 2015 onward.

Here, we report results from an effort to model the total winter abundance time series and compute TYC values following suggestions in the Red List Guidelines (IUCN, 2022) and Akçakaya *et al*. (2021). We modeled total winter abundance using four candidate models that varied in their capacity to accommodate variable change rates. We evaluated the four models for their balance between parsimony and fit using information criteria, produced Bayesian posterior distributions for TYC values from all models, and calculated model-weighted TYC summaries that incorporated uncertainty in both parameter estimation and model selection. Finally, we used the best-performing density-dependent state-space model to estimate quasi-extinction probabilities over the next 20 years to demonstrate that model choice has implications for both TYC estimates and extinction predictions.

## Methods

The data for monarch butterflies wintering in Mexico (Eastern migrants) from 1994—2018 came from Thogmartin *et al*. (2020). Data for 1993 and 2019—2021 were attained from the Monarch Joint Venture (MJV, 2022). Eastern migrant data were reported as mean hectares occupied. We converted hectares occupied to number of monarchs by multiplying hectares by 21.1 million monarchs per hectare, as was done in Walker *et al*. (2022, Thogmartin *et al*., 2017a). The data for monarch butterflies wintering along the west coast of the USA (Western migrants) from 1997—2021 were attained from the Xerces Society Western Monarch Count program (XCWMC, 2022). Estimated abundances from 1997—2021 were then summed across Eastern and Western migrants. Abundance measures from 1993—1996 for Western migrants were not readily available. Therefore, total abundances were estimated using the mean annual proportion of Western to Eastern abundance (0.0035) to avoid systematic underestimation of monarch abundance early in the time series. Note, by the proportion, that the Western subpopulation is a very small fraction of the monarch total. This effort yielded a dataset that was not identical to that used in the IUCN assessment; however, we expect that any small differences introduced at this stage had little impact on the qualitative conclusions of this analysis.

We modeled the natural log of total annual winter monarch abundance as a function of year using four candidate models. Log-transformed abundance and modeling techniques using normal distributions were employed instead of untransformed abundance and modeling techniques using Poisson or negative binomial distributions to be consistent with the monarch literature (Semmes *et al*., 2016; Schultz *et al*., 2017; Thogmartin *et al*., 2020; Walker *et al*., 2022). The candidate model set included a constant change model (Walker *et al*., 2022; hereafter “EXP”), a segmented change model (Thogmartin *et al*., 2020; hereafter “SEG”), a generalized additive model (hereafter “GAM”), and a state space model (Semmens *et al*., 2016; Schultz *et al*., 2017; hereafter “SSM”). The EXP model took the form log(*Y*_*t*_) = *α* + *β year*_*t*_ + *ϵ*_*t*_, with an intercept, a single constant change rate (here and elsewhere, exponential due to log[*Y*_*t*_]), and normally distributed errors (here and elsewhere, lognormal due to log[*Y*_*t*_]). The SEG model took the form log(*Y*_*t*_) = *α* + *β year*_*t*_ + *β*_2_ (*year*_*t*_ − *δ*) + *ϵ*_*t*_, with an intercept, an initial change rate, a second change rate after a change point, and normally distributed errors (Lunn *et al*., 2013). The GAM was a P-spline with the form log(*Y*_*t*_) = *α* + Σ_1−*K*_*β*_*k*_*B*_*k*_(*year*_*t*_) + *ϵ*_*t*_, which included an intercept, a linear combination of 3^rd^ degree B-splines with *K* = 6 knots (one per roughly five years; Smith & Edwards, 2021), and normally distributed errors (Congdon, 2014). The SSM was a stochastic Gompertz state space model that comprised a first-order autoregressive state or process component, log(*N*_*t*_) = log(*N*_*t*−1_) + *β*_0_ + *β*_1_log(*N*_*t*−1_) + *ϵ*_*t*_, with a baseline change rate, a density-dependent change-rate adjustment, and normally distributed process variation, and an observation component, log(*Y*_*t*_) = log(*N*_*t*_) + *ϵ*_*t*_, with normally distributed observation errors (Dennis *et al*., 2006; Auger-Méthé *et al*., 2021). A stochastic Gompertz state space model was chosen over a density independent state space model (Semmens *et al*., 2016; Schultz *et al*., 2017) because monarch population growth rates appear to be density dependent (Marini & Zalucki, 2017) and because density independent growth is a special case of the model (e.g., *β*_1_ = 0, Dennis *et al*., 2006). During early stages of the analysis, we took advantage of this fact and compared the density dependent model to a density independent model (*β*_1_ was forced to 0) and found that the density dependent version performed much better, based on the model selection metrics described below.

Models were analyzed in a Bayesian framework and were estimated using JAGS MCMC software (Plummer, 2017) interfaced with R statistical computing software (R Core Team, 2022). Exact specification of model structure and prior distributions can be seen in the computing code in the Supporting Information. Briefly, prior distributions for model parameters were normal or uniform, with parameter values generally considered vague (Lunn *et al*., 2013). One exception was the segmented model, where an informed prior from Thogmartin *et al*. (2020) was set for the change point parameter to promote parameter identification and MCMC convergence (Lunn *et al*., 2013). A second exception was the SSM, where we used an informed prior for sampling error from Semmens *et al*. (2016), a common approach for promoting parameter identification and MCMC convergence for state space models (Auger-Méthé *et al*., 2016). While estimating model parameters, we also computed TYC per MCMC iteration with the derived parameter *TYC* = (*N*_2021_ – *N*_2011_) / *N*_2011_ × 100. Gelman-Rubin and effective sample size metrics were used to check for adequate MCMC convergence and sample independence, respectively (Lunn *et al*., 2013), and 20,000 posterior samples were used for inference. After model estimation, we judged the general fit of the four candidate models using the Pearson correlation between observed and predicted abundances. We used the *loo* package (Vehtari *et al*., 2021) for R to compute the leave-one-out cross-validation information criterion for each model (Vehtari *et al*., 2017; hereafter “LOOIC”) and to produce model weights for each model using Stacking and Bayesian Model Averaging techniques (Yao *et al*., 2018). Pointwise likelihoods used for producing model LOOIC and weights did not include calculations for the first three observations in the time series to give time for the variance estimates from the SSM to expand from initial values and stabilize.

We found that the SSM had a much lower LOOIC than the other models, so we used it to predict monarch abundances over the next 20 years. Future abundances were computed using the state component of the SSM so that process variation, and not observation error, was incorporated into model projections. We used the posterior distribution for predicted abundance in 2041 to determine probabilities associated with monarchs dropping below quasi-extinction thresholds of 200,000, 1 million, 3 million, and 5 million individuals, which were comparable to those used in Semmens *et al*. (2016) after multiplying their four critical area thresholds by 21.1 million monarchs per hectare (Thogmartin *et al*., 2017a). We note that this approach to exploring extinction probabilities is relatively simple and phenomenological compared to more complex and mechanistic population viability analyses that employ matrix population models and stage-specific demographic rates (Morris & Doak, 2002). A benefit of this simple approach is that it requires relatively few data inputs and does not rely on explicit assumptions about matrix model structure. Nevertheless, this simple approach carries implicit assumptions that multiple factors controlling monarch populations are adequately represented by the process variation component of the state space model, and that there is little net change in these factors over the next 20 years. These are strong assumptions in a rapidly changing world. We suggest that readers not view results from this quasi-extinction analysis as robust estimates of future extinction risk, but rather as a demonstration of how predictions change considerably as new data are collected and different prediction models are employed.

## Results

The four candidate models fit the data to varying degrees (Figure 2). EXP was the poorest fitting model, with a Pearson coefficient of *r* = 0.64 for the correlation between observed and predicted abundance. The EXP model, with its single exponential change term, tended to overpredict abundance from 2011—2014 and underpredict abundance from 2015—2021 (Figure 2). The SEG and GAM models fit the data better than the EXP model, with correlation coefficients of *r* = 0.71 and *r* = 0.75, respectively (Figure 2). Both models captured the apparent shift in abundance trend predicted in Thogmartin *et al*. (2017b) and documented in Thogmartin *et al*. (2020). The GAM identified a shift around 2011 or 2012, whereas the SEG model identified a shift around 2013 or 2014 (Figure 2). The best-fitting model was the SSM, which predicted a general pattern similar to the GAM and SEG models but had a considerably tighter fit (*r* = 0.97) due to its greater flexibility (Figure 2). The posterior distribution for observation error, on the standard deviation scale, had a mean of 0.39, a standard deviation of 0.17, and a 95% credible interval of 0.01 to 0.68. The posterior distribution for process variation, on the standard deviation scale, had a mean of 0.49, a standard deviation of 0.18, and a 95% credible interval of 0.13 to 0.82. The posterior median value for the density dependence parameter was −0.30; the probability of a value less than zero, indicating negative density dependence, was 0.96, confirming results from Marini & Zalucki (2017).

**Figure 2.**
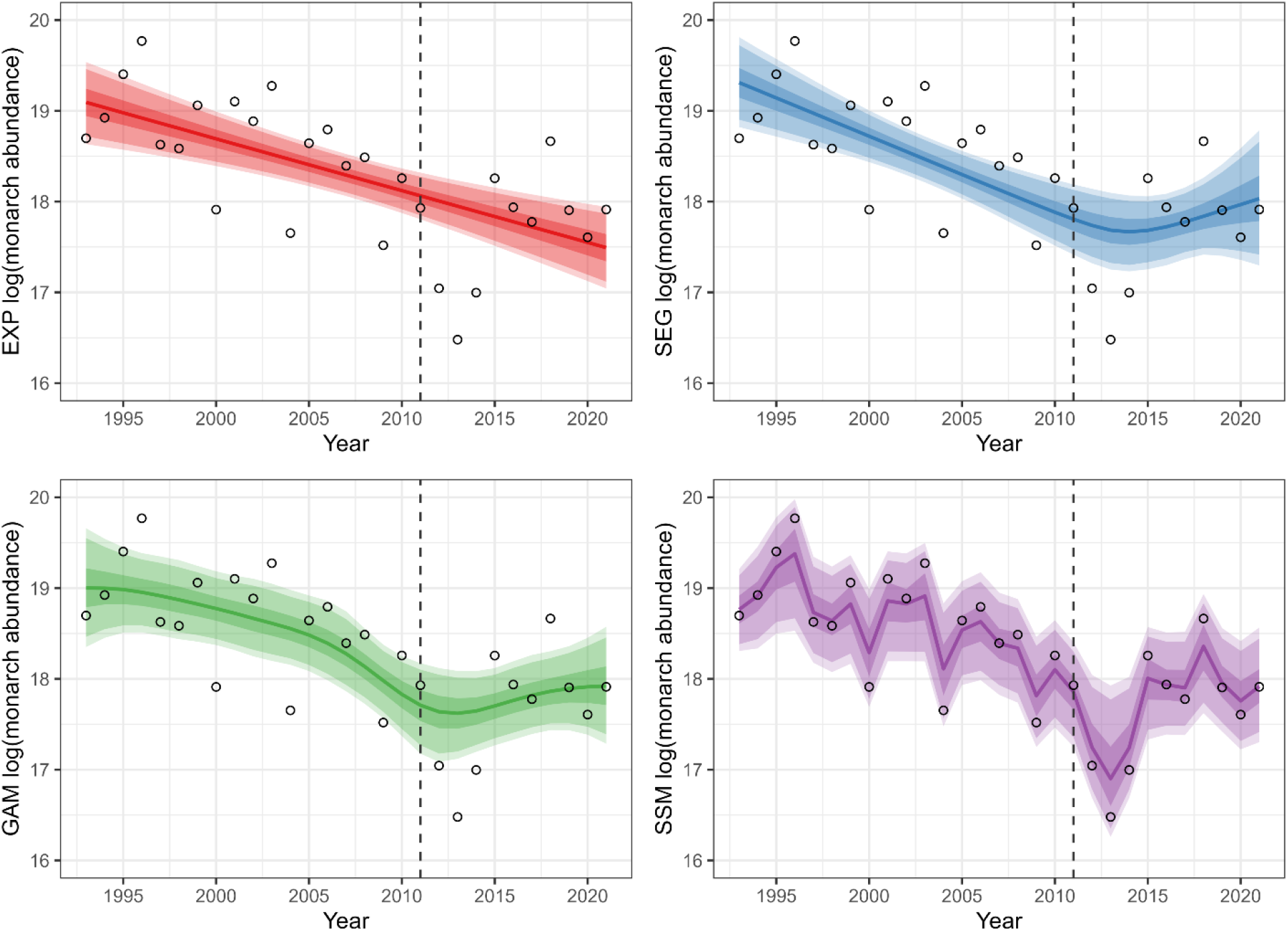
Natural log of estimated total winter abundance of migratory monarch butterflies from 1993 through 2021. Also shown are predictions from the EXP (red), SEG (blue), GAM (green), and SSM (purple) models. Model predictions are displayed as posterior medians and 50%, 90% and 95% credible intervals. The dashed vertical line in each panel shows the ten-year cutoff for estimating ten-year change values for IUCN Red List assessments.

The variation in model fit mirrored model complexity, with the simple EXP model fitting least well, the intermediate-complexity SEG and GAM fitting better, and the high-complexity SSM fitting best. We used LOOIC to judge if the increase in model complexity was justified and found that the LOOIC-best model (i.e., lowest LOOIC value) was the SSM (LOOIC = 43.89), followed by SEG (49.27), GAM (50.04), and EXP (52.29) models (Figure 3). We used Stacking and Bayesian Model Averaging techniques to create model weights (Vehtari *et al*., 2017) and found that both techniques produced model weights that were highest for the SSM (1, 0.80), intermediate for GAM (0, 0.05) and SEG (0, 0.10) models, and lowest for the EXP (0, 0.05) model. Averaged across the two methods, the model weights for SSM, GAM, SEG, and EXP models were 0.90, 0.03, 0.05, and 0.02.

**Figure 3.**
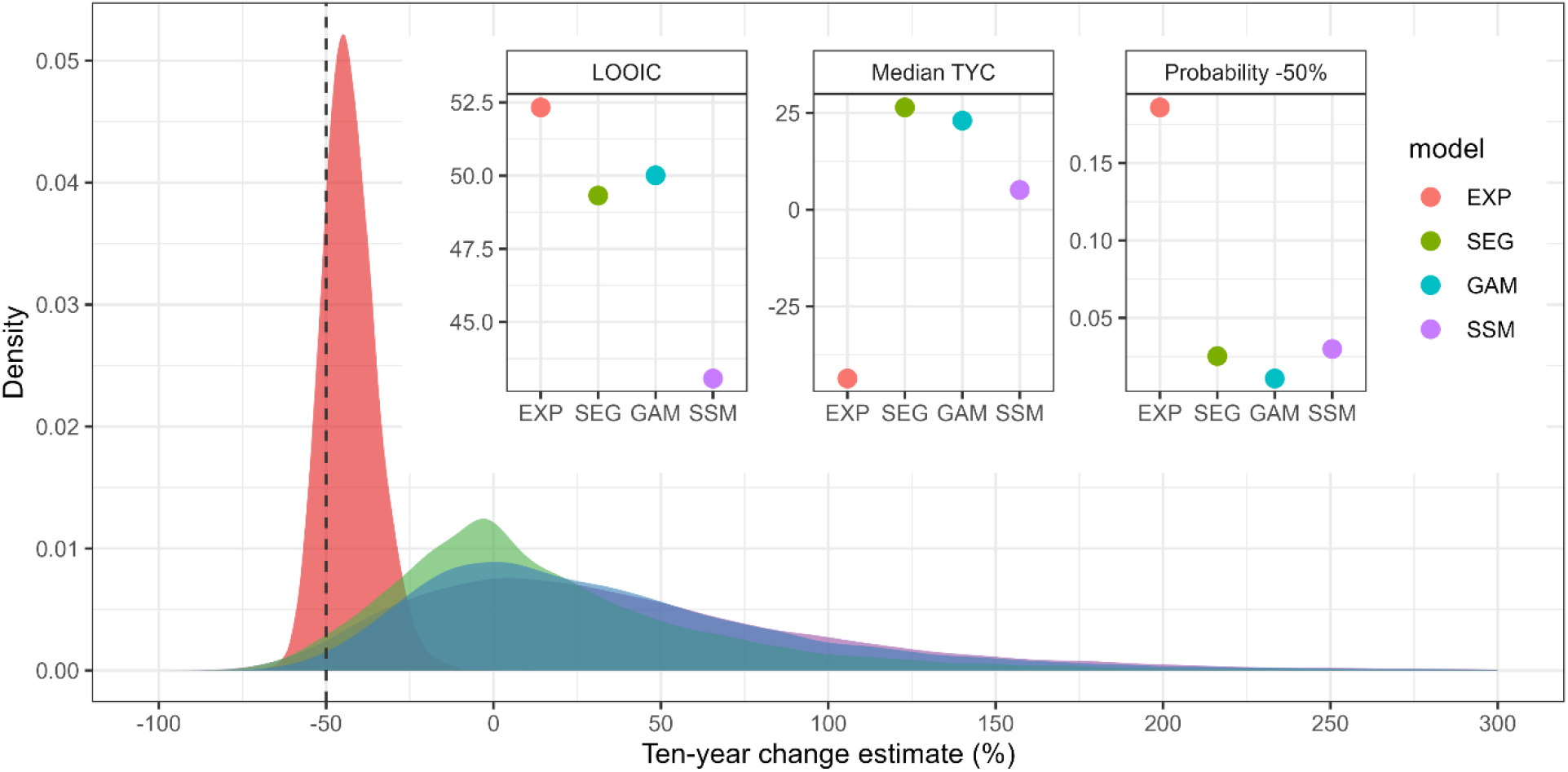
Bayesian posterior probability distributions for ten-year change (TYC) values from four candidate models. Dashed vertical line indicates a TYC value of −50%, the critical value for IUCN Red List A2ab Endangered status. Insets show the LOOIC values of the four models, where lower values indicate better predictive performance, the median TYC values from the four models, and the estimated probabilities of TYC ≤ −50% derived from the four distributions.

TYC posterior distributions for the four models clustered into two groups (Figure 3). SSM, GAM, and SEG models produced similar posterior distributions for TYC with most of the density favoring positive abundance change over the past 10 years, with posterior median TYC values of +5.34%, +22.49% and +24.90%, respectively (Figure 3). EXP was the only model favoring negative abundance change, with a posterior median value of −43.64% (Figure 3). We used the TYC posterior distributions to estimate exceedance probabilities and found that the probabilities of TYC values ≤ −50% were low for all models, at 0.03, 0.01, 0.02, and 0.19 for SSM, GAM, SEG, and EXP models, respectively (Figure 3). The probabilities of TYC values ≤ – 30% from individual models were higher, at 0.13, 0.08, 0.10, and 0.94 for SSM, GAM, SEG, and EXP models, respectively. It is possible to weight posterior summaries by the average model weights given above, yielding TYC values that consider uncertainty in both parameter estimation and model selection. When that was done, we found that the most likely TYC value for migratory monarch butterflies was +5.23%. The probability of a positive TYC was 0.56. The probability of a negative TYC value was 0.44. The probability of a TYC value at or below −30% was 0.15. Finally, the probability of a TYC value at or below −50% was 0.03.

Given that model weights for the SSM were considerably higher than those of competing models, we projected the SSM out 20 years to explore quasi-extinction probabilities. Figure 4 shows the posterior probability distribution from the SSM for the natural log of monarch abundance in 2041. The median of that distribution was exp(18.13) = 74.71 million monarch butterflies in 2041. The four vertical lines in Figure 4 highlight 2041 monarch abundances of 200,000 (dashed line, darkest gray), 1 million, 3 million, and 5 million (dashed line, lightest gray), roughly corresponding with potential quasi-extinction thresholds in Semmens *et al*. (2016). Calculating the area under the curve to the left of a given line gives the probability of a value less than or equal to the corresponding abundance in 20 years. This exercise gave probabilities of 0.008, 0.013, 0.020, and 0.026 for reaching the most conservative (200,000 monarchs) to the most liberal (5 million monarchs) quasi-extinction thresholds from Semmens *et al*. (2016). Even the most liberal quasi-extinction threshold that we have seen published, 12.8 million monarchs (Voorhies *et al*., 2019), had a very low exceedance probability of 0.055.

**Figure 4.**
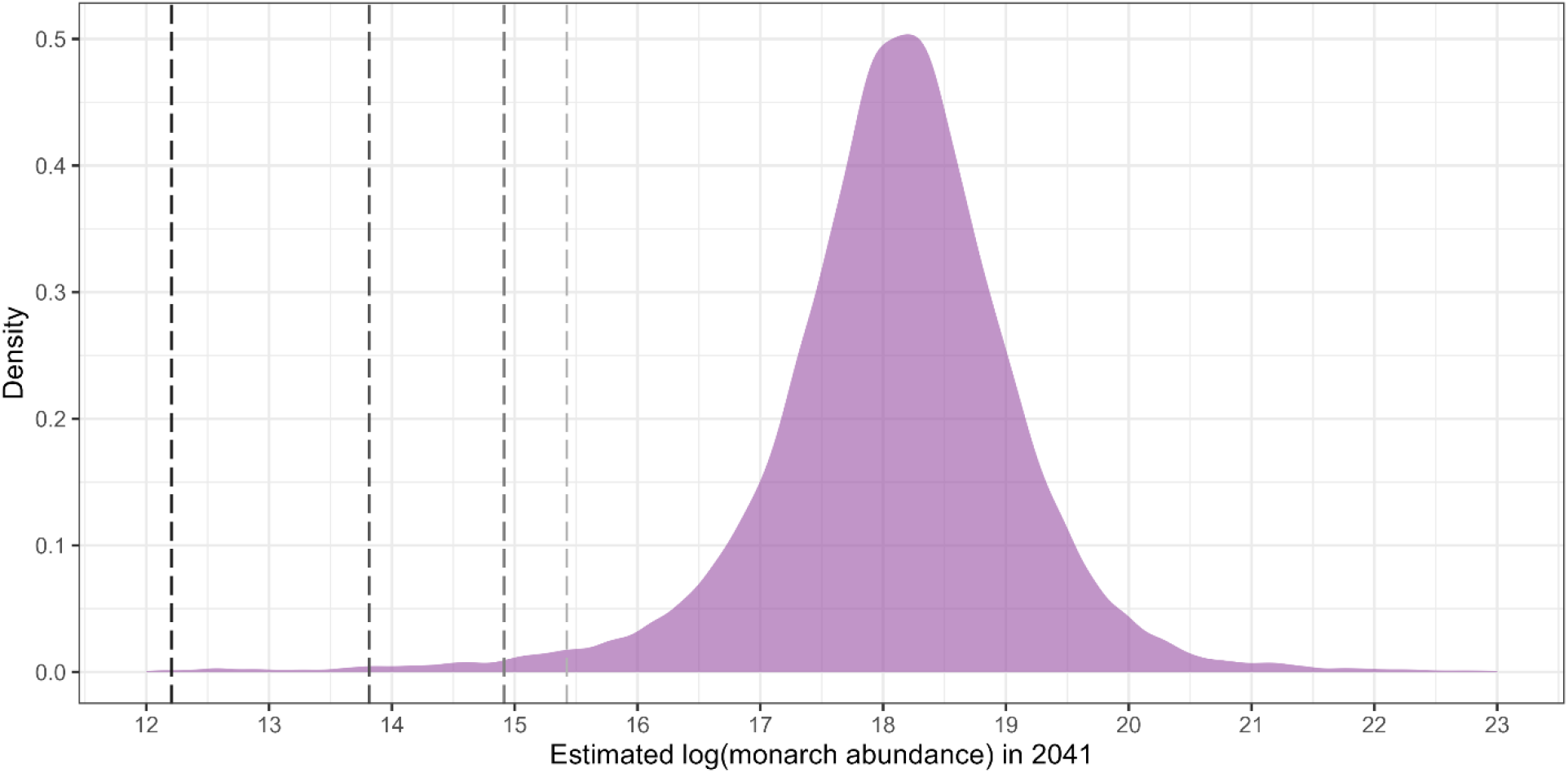
Posterior probability distribution for the natural log of monarch butterfly abundance in the year 2041 predicted from the SSM. The median of the distribution is 18.13 which translates to a population size of 74.78 million monarchs. Vertical lines represent quasi-extinction abundance thresholds comparable to Semmens *et al*. (2016), ranging from 200,000 (darkest gray dashed line) to 1 million, 3 million, and 5 million (lightest gray dashed line).

## Discussion

The IUCN Red List assessment for the migratory monarch butterfly relied heavily on an analysis of a time series of total annual winter abundance between 1993 and 2020 (Walker *et al*., 2022). That analysis yielded the conclusion that total winter monarch abundance had changed by −46% to −72% over the past ten years. These two values were considerably higher than the threshold of −30% that triggers A2ab Vulnerable status and straddled the threshold of −50% that triggers A2ab Endangered status. Consequently, the species is now on the IUCN Red List with A2ab Endangered status.

Our analysis of a nearly identical dataset yielded very different conclusions, with possible practical implications. This analysis suggested that the most likely change in abundance over the past ten years was +5.23% and that value was not statistically distinguishable from 0%, suggesting no systematic change in abundance over the past ten years (Figure 3). It also suggested that the probability of a positive TYC value (0.56) was higher than that of a negative TYC value (0.44), that the chance of TYC value below −30% was about 1 in 7, while that of a TYC value below −50% was about 1 in 33. Based on the data and models used here, the TYC values of −46% and −72% reported in the species assessment (Walker *et al*., 2022) seem highly unlikely.

Why are the conclusions of this analysis so different from those in the monarch assessment? It is not due to data source (essentially the same dataset) and minimally due to data inclusion (2021 added in this analysis), preparation (1993—1996 estimated for Western migrants in this analysis), and computing procedures. This is evident when comparing the posterior median TYC value from the EXP model, −43.64%, to the analogous value from the monarch assessment, −46%. Rather, differences are mostly due to which models were fit to the full time series. In the monarch species assessment, two constant-change models were fit to the data, two point-estimates for TYC were computed, and those two TYC point estimates were pooled with a third point estimate from the literature to represent uncertainty in population change over the last ten years. That approach was not incorrect but could be considered incomplete.

In this analysis we took extra steps intended to improve the quality of TYC estimates for the monarch butterfly. The additional steps taken were those recommended in the Red List Guidelines (IUCN, 2022) and Akçakaya *et al*., (2021) specifically for invertebrates. The first addition was to evaluate the assumption of a constant population change rate and to report TYC values derived from models that accommodate variable change rates if the assumption was not supported. Previous work (Thogmartin *et al*., 2020) and this analysis both demonstrate that a constant rate assumption is not appropriate for this time series. Therefore, TYC values from constant rate models should not be heavily weighted in a monarch assessment. Another addition was the detailed treatment of uncertainty in TYC values. In this analysis we used Bayesian inference methods that produced posterior distributions for TYC. We used posterior distributions from the four candidate models, along with model weights, to produce posterior summaries that accounted for uncertainty in both parameter estimation and model selection. These posteriors allowed probabilities to be assessed for any TYC value (Supporting Information). While Bayesian uncertainty analysis is not specifically recommended, other forms of probabilistic evaluation, such as Monte Carlo methods, are recommended in Red List Guidelines (IUCN, 2022).

While we have focused mostly on the statistical dimensions of model selection during species assessment, Red List Guidelines (IUCN, 2022) also ask assessors to choose candidate models based on putative mechanisms of population decline. In cases where a species has lost a large area of historic habitat, but that loss has abated significantly, Akçakaya *et al*. (2021) recommend against using constant rate models to calculate TYC. This scenario resembles that of monarch butterflies breeding in agricultural landscapes of the Midwestern USA, which experienced a large loss of habitat between 1990 and 2010 as the proportion of corn and soybean fields with herbicide applied increased from near 0% to near 90% (Thogmartin *et al*., 2017b). As herbicide use and habitat loss stabilized, Thogmartin *et al*. (2017b) concluded that further herbicide-associated declines in monarchs were not expected, and that monarchs would likely level off at a new stationary population size. This prediction seems well supported by results from both Thogmartin *et al*. (2020) and this analysis (Figure 2) and underscores the conclusion that constant change models may not be the best choice for modeling monarch butterfly populations.

Total winter abundance projections for 2041 from the best-performing, density-dependent, state space model gave chances of quasi-extinction over the next 20 years between 0.8% and 2.6% for abundance thresholds ranging from 200,000 to 5 million monarchs, respectively (Figure 4). These chances were an order of magnitude lower than previous estimates that ranged from 11% to 53% in Semmens *et al*. (2016) and from 5% to 43% in the monarch assessment (Walker *et al*., 2022). (Note that the threshold for Red List Criterion E Endangered status is 20%; IUCN, 2022). As mentioned above, the intention of our quasi-extinction analysis was not to produce the latest round of robust extinction predictions. Rather, the results reported here were intended to demonstrate how incorporating recent data and selecting different prediction methods affects extinction predictions. The quasi-extinction probabilities from Semmens *et al*. (2016) were derived from a state-space model that included abundance data through 2014, right around the time when monarch declines were potentially easing (Thogmartin *et al*., 2020; this study). Given the data available at that time, relatively high estimates of extinction probabilities are not surprising. Another recent development is the clear demonstration of density dependence in multiple time series of Monarch abundances (Marini & Zalucki, 2017; this study). The quasi-extinction probabilities from Semmens *et al*. (2016) were derived from methods that assumed density independence. It is common that methods assuming density independence will produce relatively high extinction probabilities compared to methods assuming density dependence, because density dependent growth rates allow populations to rebound quickly following years of low abundance, with strong implications for population persistence (Morris & Doak, 2002). A clear empirical example of the monarch’s ability to rebound is seen in winter abundance data from the West Coast of the USA (Supporting Information), where abundance in 2021 (247,246 monarchs) was around 130 times higher than that in the preceding year (2020; 1,899 monarchs). Under several simplifying assumptions, that equates with a finite rate of increase of 247,246 / 1,899 = 130.20 or an instantaneous rate of increase of log(130.20) = 4.87.

In summary, it appears that the conclusions of the current IUCN assessment for the monarch butterfly are derived from an incomplete evaluation of abundance changes over the past ten years. We suggest that a more thorough analysis of recent overwintering abundances will lead to an improved IUCN assessment for monarch butterflies. We recommend that other researchers evaluating monarch conservation status consider using models with variable change rates, as models with constant change rates may not accurately predict the trajectory of monarch abundances into the future.

## Acknowledgements

We thank three colleagues for helpful comments on a previous draft of this manuscript. MSC acknowledges funding from USDA Hatch #DEL00774.

## References

Akçakaya, H.R., Hochkirch, A., Bried, J.T., van Grunsven, R.H., Simaika, J.P., De Knijf, G. & Henriques, S. (2021) Calculating population reductions of invertebrate species for IUCN Red List assessments. Journal of Insect Conservation, 25 377–382.

Auger-Méthé, M., Field, C., Albertsen, C.M., Derocher, A.E., Lewis, M.A., Jonsen, I.D. & Mills Flemming, J. (2016) State-space models’ dirty little secrets: even simple linear Gaussian models can have estimation problems. Scientific Reports, 6, 26677.

Auger-Méthé, M., Newman, K., Cole, D., Empacher, F., Gryba, R., King, A.A., Leos-Barajas, V., Mills Flemming, J., Nielsen, A., Petris, G. & Thomas, L. (2021) A guide to state–space modeling of ecological time series. Ecological Monographs 91, e01470.

Congdon, P. (2014) Applied Bayesian modelling. John Wiley & Sons, West Sussex, UK.

Dennis, B., Ponciano, J.M., Lele, S.R., Taper, M.L. & Staples, D.F. (2006) Estimating density dependence, process noise, and observation error. Ecological Monographs, 76, 323–341.

Fewster, R.M., Buckland, S.T., Siriwardena, G.M., Baillie, S.R. & Wilson, J.D. (2000) Analysis of population trends for farmland birds using generalized additive models. Ecology, 81, 1970–1984.

Flockhart, D.T.T., Martin T.G. & Norris D.R. (2012) Experimental examination of intraspecific density-dependent competition during the breeding period in monarch butterflies (*Danaus plexippus*). PLoS ONE, 7, e45080.

Hooten, M.B. & Hobbs, N.T. (2015) A guide to Bayesian model selection for ecologists. Ecological Monographs, 85, 3–28.

IUCN Standards and Petitions Committee (2022) Guidelines for Using the IUCN Red List Categories and Criteria. Version 15.1. Prepared by the Standards and Petitions Committee. <https://www.iucnredlist.org/documents/RedListGuidelines.pdf> 28th December 2022.

Lunn, D., Jackson, C., Best, N., Thomas, A. & Spiegelhalter, D. (2013) The BUGS Book. A Practical Introduction to Bayesian Analysis. Chapman Hall, London, UK.

Marini, L. & Zalucki, M.P. (2017) Density-dependence in the declining population of the monarch butterfly. Scientific Reports, 7, 13957.

Monarch Joint Venture (MJV) (2022) Eastern monarch population holds steady at 2.84 hectares. <https://monarchjointventure.org/blog/eastern-monarchs-hold-steady> 28th December 2022.

Morris, W.F. & Doak, D.F. (2002) Quantitative Conservation Biology. Sinauer, Sunderland, Massachusetts, USA.

Plummer, M. (2017) JAGS Version 4.3.0 user manual. <https://sourceforge.net/projects/mcmc-jags/files/Manuals/4.x/jags_user_manual.pdf/download> 28th December 2022.

R Core Team (2022). R: A language and environment for statistical computing. R Foundation for Statistical Computing, Vienna, Austria. <https://www.R-project.org/>.

Schultz, C.B., Brown, L.M., Pelton, E. & Crone, E.E. (2017) Citizen science monitoring demonstrates dramatic declines of monarch butterflies in western North America. Biological Conservation, 214, 343–346.

Semmens, B.X., Semmens, D.J., Thogmartin, W.E., Wiederholt, R., López-Hoffman, L., Diffendorfer, J.E., Pleasants, J.M., Oberhauser, K.S. & Taylor, O.R. (2016) Quasi-extinction risk and population targets for the Eastern migratory population of monarch butterflies (*Danaus plexippus*). Scientific Reports, 6, 23265.

Smith, A.C. & Edwards, B.P. (2021) North American Breeding Bird Survey status and trend estimates to inform a wide range of conservation needs, using a flexible Bayesian hierarchical generalized additive model. The Condor, 123, duaa065.

Thogmartin, W.E., Diffendorfer, J.E., López-Hoffman, L., Oberhauser, K., Pleasants, J., Semmens, B.X., Semmens, D., Taylor, O.R. & Wiederholt, R. (2017a) Density estimates of monarch butterflies overwintering in central Mexico. PeerJ, 5, p.e3221.

Thogmartin, W.E., Wiederholt, R., Oberhauser, K., Drum, R.G., Diffendorfer, J.E., Altizer, S., Taylor, O.R., Pleasants, J., Semmens, D., Semmens, B. & Erickson, R. (2017b) Monarch butterfly population decline in North America: identifying the threatening processes. Royal Society Open Science, 4, 170760.

Thogmartin, W.E., Szymanski, J.A. & Weiser, E.L. (2020) Evidence for a growing population of Eastern migratory monarch butterflies is currently insufficient. Frontiers in Ecology and Evolution, 8, 43.

Vehtari, A., Gelman, A. & Gabry, J. (2017) Practical Bayesian model evaluation using leave-one-out cross-validation and WAIC. Statistics and Computing, 27, 1413–1432.

Vehtari, A., Gelman, A., Gabry, J. & Yao, Y. (2021) Package ‘loo’. Efficient Leave-One-Out Cross-Validation and WAIC for Bayesian Models. <http://mc-stan.org/loo/index.html> 28th December 2022.

Voorhies, K.J., Szymanski, J., Nail, K.R. & Fidino, M. (2019) A method to project future impacts from threats and conservation on the probability of extinction for North American migratory monarch (*Danaus plexippus*) populations. Frontiers in Ecology and Evolution, 7, 384.

Walker, A., Oberhauser, K.S., Pelton, E.M., Pleasants, J.M. & Thogmartin, W.E. (2022) *Danaus plexippus* ssp. *plexippus* (errata version published in 2022). The IUCN Red List of Threatened Species 2022: e.T194052138A219151401. <https://dx.doi.org/10.2305/IUCN.UK.2022-1.RLTS.T194052138A219151401.en> 28th December 2022.

Xerces Society Western Monarch Count (XSWMC) (2022) Western Monarch Thanksgiving Count and New Year’s Count Data, 1997–2021. <http://www.westernmonarchcount.org> 28th December 2022.

Yao, Y., Vehtari, A., Simpson, D. & Gelman, A. (2018) Using stacking to average Bayesian predictive distributions (with discussion). Bayesian Analysis, 13, 917–1007.

