## Supporting Information for "Change in Monarch Butterfly Winter Abundance Over the Past Decade: A Red List Perspective"

**PREPRINT SUPPORTING INFORMATION**

Timothy D. Meehan

*National Audubon Society, New York City, New York USA,*

Michael S. Crossley

*University of Delaware, Newark, Delaware USA,*

### Note to Readers

This work has undergone friendly review from colleagues but has not yet undergone rigorous, anonymous peer review. That process is now underway, and the manuscript will be updated when it is complete.

**Supporting Information A.** Table of model-weighted percentiles from the pooled posterior for ten-year change (TYC), which can be used to approximate probabilities of different TYC values. For example, line 3, columns 1 and 2, show that the probability of TYC ≤ -50 is approximately 3%. Line 14, columns 1 and 2, show that the probability of TYC ≤ -30 is approximately 14%. Line 20, columns 3 and 4, show that the probability of TYC ≤ 0 is approximately 44%. Line 1, columns 5 and 6, show that the model weighted posterior median TYC is approximately 5.23%.

| TYC | Percentile | TYC | Percentile | TYC | Percentile | TYC | Percentile |
|---|---|---|---|---|---|---|---|
| -59.51 | 1% | -17.30 | 25% | **5.23** | **50%** | 39.14 | 75% |
| -54.88 | 2% | -16.22 | 26% | 6.25 | 51% | 41.26 | 76% |
| **-50.85** | **3%** | -15.21 | 27% | 7.27 | 52% | 43.28 | 77% |
| -48.09 | 4% | -14.30 | 28% | 8.24 | 53% | 45.57 | 78% |
| -45.88 | 5% | -13.16 | 29% | 9.43 | 54% | 47.76 | 79% |
| -43.95 | 6% | -12.21 | 30% | 10.59 | 55% | 50.45 | 80% |
| -42.04 | 7% | -11.17 | 31% | 11.58 | 56% | 53.13 | 81% |
| -40.25 | 8% | -10.25 | 32% | 12.84 | 57% | 55.73 | 82% |
| -38.54 | 9% | -9.32 | 33% | 14.07 | 58% | 58.80 | 83% |
| -36.91 | 10% | -8.44 | 34% | 15.22 | 59% | 61.98 | 84% |
| -35.17 | 11% | -7.56 | 35% | 16.50 | 60% | 65.30 | 85% |
| -33.69 | 12% | -6.59 | 36% | 17.85 | 61% | 68.37 | 86% |
| -32.29 | 13% | -5.52 | 37% | 19.04 | 62% | 71.79 | 87% |
| **-30.73** | **14%** | -4.72 | 38% | 20.27 | 63% | 75.60 | 88% |
| -29.29 | 15% | -4.05 | 39% | 21.56 | 64% | 80.26 | 89% |
| -28.13 | 16% | -3.24 | 40% | 22.95 | 65% | 84.98 | 90% |
| -26.88 | 17% | -2.48 | 41% | 24.35 | 66% | 90.38 | 91% |
| -25.44 | 18% | -1.72 | 42% | 25.70 | 67% | 95.74 | 92% |
| -24.13 | 19% | -0.97 | 43% | 27.04 | 68% | 102.05 | 93% |
| -23.05 | 20% | **-0.23** | **44%** | 28.62 | 69% | 110.22 | 94% |
| -21.82 | 21% | 0.57 | 45% | 30.20 | 70% | 119.10 | 95% |
| -20.76 | 22% | 1.47 | 46% | 32.04 | 71% | 130.87 | 96% |
| -19.66 | 23% | 2.38 | 47% | 33.64 | 72% | 146.59 | 97% |
| -18.45 | 24% | 3.33 | 48% | 35.31 | 73% | 169.35 | 98% |
| | | 4.27 | 49% | 37.39 | 74% | 210.28 | 99% |

**Supporting Information B**. All data and R and JAGS computing code used for this analysis.

```
# setup ------------------------------------------------------------------------
library(scales)
library(Hmisc)
library(cowplot)
library(ggplot2)
library(splines)
library(loo)
library(MCMCvis)
library(R2jags)
library(dplyr)
setwd(getwd())
theme_set(theme_bw())
# ----------------------------------------------------------------------------


# get data ---------------------------------------------------------------------
md0 <- data.frame(
  pop="Eastern migrants",
  year = 1993:2021,
  index = c(6.23, 7.81,12.61,18.19,5.77,5.56,8.97,2.83,9.36,7.54,11.12,
            2.19,5.91,6.87,4.61,5.06,1.92,4.02,2.89,1.19,0.67,1.13,4.01,
            2.91,2.48,6.05, 2.83, 2.10, 2.84)
) %>%
  mutate(index=round(index * 21100000,0)) %>%
  bind_rows(data.frame(
    pop="Western migrants",
    year = 1997:2021,
    index = c(1235490,564349,267574,390057,209570,99353,254378,205085,
              218679,221058,86437,131889,58468,143204,222525,144812,
              211275,234731,292888,298464,192624,27721,29436,1899,
              247246)))

# sum across regional populations
md1 <- md0 %>%
  group_by(year) %>%
  summarise(sites=n(),
            west_prop=min(index) / max(index),
            index=sum(index, na.rm=F))

# correct first four years using mean proportion from west
cf <- md1 %>% filter(west_prop!=1) %>% pull(west_prop) %>% mean() + 1
md1$index[md1$west_prop==1] <- md1$index[md1$west_prop==1] * cf
md1 <- md1 %>% dplyr::select(year, index) %>%
  mutate(yr=year-min(year)+1, log_idx=log(index))

# plot data
md1 %>% mutate(efit=exp(fitted(lm(log_idx~year, data=md1))),
               lfit=fitted(lm(index~year, data=md1))) %>%
  ggplot(aes(x=year, y=index)) +
  geom_line(col="gray60") +
  geom_point(pch=21, fill="white", size=1.9) +
  labs(x="Year", y="Monarch abundance") +
  scale_y_continuous(label=comma, limits=c(0, 400000000),
                     breaks=seq(0, 400000000, 50000000)) +
  scale_x_continuous(breaks=seq(1995, 2020, 5)) +
  geom_line(aes(y=efit), col="#e41a1c", lty=5) +
  geom_line(aes(y=lfit), col="#e41a1c", lty=1)
```

```
ggsave("figure_1.tiff", width = 8.66, height = 4.33, dpi = 600, units = "in")

# some other data properties
n_years <- nrow(md1)
# ------------------------------------------------------------------------------


# exponential model ------------------------------------------------------------
# write jags model
cat(file = "exp_reg.txt", "
model {
### priors
  alpha ~ dnorm(0, 0.001)
  beta ~ dnorm(0, 0.001)
  sigma ~ dunif(0, 10)
### likelihood
  for (i in 1:n) {
    y[i] ~ dnorm(mu[i], tau)
    mu[i] <- alpha + beta*x[i]
  }
### derived
  tau <- 1/ (sigma * sigma)
  tyc <- ((exp(mu[29]) - exp(mu[19])) / exp(mu[19])) * 100
  for(i in 4:n){
    loglik[i] <- logdensity.norm(y[i], mu[i], tau)
  }
}
")

# bundle jags data
dat_exp <- list(x = md1$yr, y = md1$log_idx, n = n_years)

# parameters to monitor
params <- c("alpha","beta", "sigma", "tyc", "loglik", "mu")

# mcmc settings
ni <- 15000000; nt <- 1000; nb <- 5000000; nc <- 2; nad <- 10000

# call jags and summarise posteriors
exp_out <- jags(dat_exp, inits=NULL, params, "exp_reg.txt",
                n.thin=nt, n.chains=nc, n.burnin=nb, n.iter=ni)

# check some diagnostics
(exp_sum <- MCMCsummary(exp_out))
MCMCtrace(exp_out, "alpha", pdf=F)
MCMCtrace(exp_out, "beta", pdf=F)
MCMCtrace(exp_out, "sigma", pdf=F)
MCMCtrace(exp_out, "tyc", ISB=T, pdf=F)
MCMCtrace(exp_out, "mu[5]", ISB=F, pdf=F)
MCMCtrace(exp_out, "mu[10]", ISB=F, pdf=F)
MCMCtrace(exp_out, "mu[15]", ISB=F,pdf=F)
# ------------------------------------------------------------------------------


# segmented exponential model --------------------------------------------------
# write jags model
cat(file = "seg_reg.txt", "
model {
### priors
  alpha ~ dnorm(0, 0.001)
  for(j in 1:2){
```

```
    beta[j] ~ dunif(-1, 1)
    }
  sigma ~ dunif(0, 10)
  theta ~ dunif(15, 25)
### likelihood
  for (i in 1:n) {
    y[i] ~ dnorm(mu[i], tau)
    mu[i] <- alpha + beta[1]*x[i] + beta[2]*(x[i] - theta) * step(x[i] - theta)
  }
### derived
  tau <- 1/ (sigma * sigma)
  tyc <- ((exp(mu[29]) - exp(mu[19])) / exp(mu[19])) * 100
  for(i in 4:n){
    loglik[i] <- logdensity.norm(y[i], mu[i], tau)
  }
}
")

# parameters to monitor
params <- c("alpha","beta", "sigma", "theta", "tyc", "loglik", "mu")

# call jags and summarise posteriors
seg_out <- jags(dat_exp, inits=NULL, params, "seg_reg.txt",
                n.thin=nt, n.chains=nc, n.burnin=nb, n.iter=ni)

# check some diagnostics
(seg_sum <- MCMCsummary(seg_out))
MCMCtrace(seg_out, "alpha", pdf=F)
MCMCtrace(seg_out, "beta", pdf=F)
MCMCtrace(seg_out, "sigma", pdf=F)
MCMCtrace(seg_out, "theta", pdf=F)
MCMCtrace(seg_out, "tyc", ISB=T, pdf=F)
MCMCtrace(seg_out, "mu[5]", ISB=F, pdf=F)
MCMCtrace(seg_out, "mu[10]", ISB=F, pdf=F)
MCMCtrace(seg_out, "mu[15]", ISB=F,pdf=F)
# ------------------------------------------------------------------------------


# state space model ------------------------------------------------------------
# write stochastic gompertz state space model
cat(file = "ssm_mod.txt", "
model {
### priors
  mu[1] ~ dnorm(y[1], ((0.67-0.21)/1.97)^-2)
  b0 ~ dnorm(0, 0.001)
  b1 ~ dnorm(0, 0.001)
  tau.obs <- 1 / (sigma.obs * sigma.obs)
  sigma.obs ~ dnorm(0.44, ((0.67-0.21)/1.97)^-2)
  tau.proc <- 1 / (sigma.proc * sigma.proc)
  sigma.proc ~ dunif(0, 100)
### likelihood
  for (i in 2:n){
    y[i] ~ dnorm(mu[i], tau.obs)
    mu[i] <- mu[i - 1] + rate[i]
    rate[i] ~ dnorm(b0 + b1 * (mu[i - 1]), tau.proc)
  }
### derived
  tyc <- ((exp(mu[29]) - exp(mu[19])) / exp(mu[19])) * 100
  for(i in 4:n){
    loglik[i] <- logdensity.norm(y[i], mu[i], tau.obs)
  }
```

```
}
")

# bundle jags data
dat_ssm <- list(y = md1$log_idx, n = n_years)

# parameters to monitor
parameters <- c("b0", "b1", "tyc", "sigma.proc", "sigma.obs","rate", "loglik",
                "mu")

# call jags and summarise posteriors
ssm_out <- jags(dat_ssm, inits=NULL, parameters, "ssm_mod.txt",
                n.chains=nc, n.thin=nt, n.iter=ni, n.burnin=nb)

# check a few diagnostics
(ssm_sum <- MCMCsummary(ssm_out))
MCMCtrace(ssm_out, "b0", pdf=F)
MCMCtrace(ssm_out, "b1", pdf=F)
MCMCtrace(ssm_out, "sigma.proc", pdf=F)
MCMCtrace(ssm_out, "sigma.obs", pdf=F)
MCMCtrace(ssm_out, "tyc", ISB=T, pdf=F)
MCMCtrace(ssm_out, "mu[5]", ISB=F, pdf=F)
MCMCtrace(ssm_out, "mu[10]", ISB=F, pdf=F)
MCMCtrace(ssm_out, "mu[15]", ISB=F,pdf=F)
sum(MCMCchains(ssm_out, "b1")<0)/20000
summary(MCMCchains(ssm_out, "rate")[,1])
mean(MCMCchains(ssm_out, "rate")[,19:28])

# set rate to zero to see carrying capacity
a <- as.numeric(MCMCsummary(ssm_out, "b0")[4])
b <- as.numeric(MCMCsummary(ssm_out, "b1")[4])
c <- b + 1
nstat <- a / (1 - (b + 1))
# ---------------------------------------------------------------------------


# gam model -----------------------------------------------------------------
# write jags model
cat(file = "gam_mod.txt", "
model {
### rw prior on beta
  beta[1] ~ dnorm(0, 0.001)
  for (i in 2:n_knots) {
    beta[i] ~ dnorm(beta[i-1], sigma_b^-2)
  }
### priors on beta values
  sigma ~ dunif(0, 10)
  sigma_b ~ dunif(0, 10)
### likelihood
  for (i in 1:n) {
    y[i] ~ dnorm(mu[i], sigma^-2)
    mu[i] <- inprod(B[i, ], beta)
  }
### derived
  tyc <- ((exp(mu[29]) - exp(mu[19])) / exp(mu[19])) * 100
  for(i in 4:n){
    loglik[i] <- logdensity.norm(y[i], mu[i], sigma^-2)
  }
}
")
```

```
# create spline structure
bs_bbase <- function(x, xl = min(x), xr = max(x), nseg = 6, deg = 3) {
  dx <- (xr - xl) / nseg
  knots <- seq(xl - deg * dx, xr + deg * dx, by = dx)
  get_bs_matrix <- matrix(bs(x, knots = knots, degree = deg, Boundary.knots =
                              c(knots[1], knots[length(knots)])), nrow = length(x))
  bs_matrix <- get_bs_matrix[, -c(1:deg, ncol(get_bs_matrix):(ncol(get_bs_matrix) - deg))]
  return(bs_matrix)
}
B <- bs_bbase(md1$yr, nseg = 6)

# bundle jags data
dat_gam <- list(n = n_years, y = md1$log_idx, B = B, n_knots = ncol(B))

# parameters to monitor
parameters <- c("beta", "loglik", "mu", "tyc")

# call jags and summarise posteriors
gam_out <- jags(dat_gam, inits=NULL, parameters, "gam_mod.txt",
                n.chains=nc, n.thin=nt, n.iter=ni, n.burnin=nb)

# check a few diagnostics
(gam_sum <- MCMCsummary(gam_out))
MCMCtrace(gam_out, "tyc", ISB=T, pdf=F)
MCMCtrace(gam_out, "mu[5]", ISB=F, pdf=F)
MCMCtrace(gam_out, "mu[10]", ISB=F, pdf=F)
MCMCtrace(gam_out, "mu[15]", ISB=F,pdf=F)
MCMCtrace(gam_out, "beta[1]", ISB=F, pdf=F)
# ------------------------------------------------------------------------------


# get predictions ---------------------------------------------------------------
# compute fitted values
exp_mus <- as.data.frame(t(apply(t(exp_out$BUGSoutput$sims.list$mu), 1, quantile,
                                 probs=c(0.025, 0.05, 0.25, 0.5, 0.75, 0.95, 0.975)))) %>%
  rename_with(~ paste0("exp", gsub("%", "", .)))
seg_mus <- as.data.frame(t(apply(t(seg_out$BUGSoutput$sims.list$mu), 1, quantile,
                                 probs=c(0.025, 0.05, 0.25, 0.5, 0.75, 0.95, 0.975)))) %>%
  rename_with(~ paste0("seg", gsub("%", "", .)))
ssm_mus <- as.data.frame(t(apply(t(ssm_out$BUGSoutput$sims.list$mu), 1, quantile,
                                 probs=c(0.025, 0.05, 0.25, 0.5, 0.75, 0.95, 0.975)))) %>%
  rename_with(~ paste0("ssm", gsub("%", "", .)))
gam_mus <- as.data.frame(t(apply(t(gam_out$BUGSoutput$sims.list$mu), 1, quantile,
                                 probs=c(0.025, 0.05, 0.25, 0.5, 0.75, 0.95, 0.975)))) %>%
  rename_with(~ paste0("gam", gsub("%", "", .)))
md2 <- md1 %>% bind_cols(exp_mus) %>% bind_cols(seg_mus) %>%
  bind_cols(ssm_mus) %>% bind_cols(gam_mus) %>%
  mutate(exp_res=log_idx-exp50, seg_res=log_idx-seg50,
         ssm_res=log_idx-ssm50, gam_res=log_idx-gam50)

# get fit stats
dev_exp <- round(exp_out$BUGSoutput$mean$deviance, 2)
dev_seg <- round(seg_out$BUGSoutput$mean$deviance, 2)
dev_ssm <- round(ssm_out$BUGSoutput$mean$deviance, 2)
dev_gam <- round(gam_out$BUGSoutput$mean$deviance, 2)
cor_exp <- round(cor(exp_out$BUGSoutput$mean$mu, md1$log_idx), 2)
cor_seg <- round(cor(seg_out$BUGSoutput$mean$mu, md1$log_idx), 2)
cor_ssm <- round(cor(ssm_out$BUGSoutput$mean$mu, md1$log_idx), 2)
cor_gam <- round(cor(gam_out$BUGSoutput$mean$mu, md1$log_idx), 2)

# get logliks
```

```
exp_ll <- exp_out$BUGSoutput$sims.list$loglik
seg_ll <- seg_out$BUGSoutput$sims.list$loglik
ssm_ll <- ssm_out$BUGSoutput$sims.list$loglik
gam_ll <- gam_out$BUGSoutput$sims.list$loglik
exp_reff <- relative_eff(exp(exp_ll), chain_id = rep(1:2, each = 10000))
seg_reff <- relative_eff(exp(seg_ll), chain_id = rep(1:2, each = 10000))
ssm_reff <- relative_eff(exp(ssm_ll), chain_id = rep(1:2, each = 10000))
gam_reff <- relative_eff(exp(gam_ll), chain_id = rep(1:2, each = 10000))

# get looic
loic_exp <- round(loo(exp_ll, r_eff=exp_reff)$estimates['looic','Estimate'], 2)
loic_seg <- round(loo(seg_ll, r_eff=seg_reff)$estimates['looic','Estimate'], 2)
loic_ssm <- round(loo(ssm_ll, r_eff=ssm_reff)$estimates['looic','Estimate'], 2)
loic_gam <- round(loo(gam_ll, r_eff=gam_reff)$estimates['looic','Estimate'], 2)

# make ts plots
ts_exp <- ggplot(md2, aes(x=year))  +
  geom_ribbon(aes(ymin=exp2.5, ymax=exp97.5), alpha=0.2, fill="#e41a1c") +
  geom_ribbon(aes(ymin=exp5, ymax=exp95), alpha=0.3, fill="#e41a1c") +
  geom_ribbon(aes(ymin=exp25, ymax=exp75), alpha=0.4, fill="#e41a1c") +
  geom_line(aes(y=exp50), linewidth=0.8, col="#e41a1c") + geom_point(aes(y=log_idx), pch=1) +
  labs(x="Year", y="EXP log(monarch abundance)") +
  scale_y_continuous(limits=c(16, 20)) +
  scale_x_continuous(breaks=seq(1995, 2020, 5)) +
  geom_vline(xintercept=2011, lty=2, col="gray20")

ts_seg <- ggplot(md2, aes(x=year))  +
  geom_ribbon(aes(ymin=seg2.5, ymax=seg97.5), alpha=0.2, fill="#377eb8") +
  geom_ribbon(aes(ymin=seg5, ymax=seg95), alpha=0.3, fill="#377eb8") +
  geom_ribbon(aes(ymin=seg25, ymax=seg75), alpha=0.4, fill="#377eb8") +
  geom_line(aes(y=seg50), linewidth=0.8, col="#377eb8") + geom_point(aes(y=log_idx), pch=1) +
  labs(x="Year", y="SEG log(monarch abundance)") +
  scale_y_continuous(limits=c(16, 20)) +
  scale_x_continuous(breaks=seq(1995, 2020, 5)) +
  geom_vline(xintercept=2011, lty=2, col="gray20")

ts_ssm <- ggplot(md2, aes(x=year))  +
  geom_ribbon(aes(ymin=ssm2.5, ymax=ssm97.5), alpha=0.2, fill="#984ea3") +
  geom_ribbon(aes(ymin=ssm5, ymax=ssm95), alpha=0.3, fill="#984ea3") +
  geom_ribbon(aes(ymin=ssm25, ymax=ssm75), alpha=0.4, fill="#984ea3") +
  geom_line(aes(y=ssm50), linewidth=0.8, col="#984ea3") + geom_point(aes(y=log_idx), pch=1) +
  labs(x="Year", y="SSM log(monarch abundance)") +
  scale_y_continuous(limits=c(16, 20)) +
  scale_x_continuous(breaks=seq(1995, 2020, 5)) +
  geom_vline(xintercept=2011, lty=2, col="gray20")

ts_gam <- ggplot(md2, aes(x=year))  +
  geom_ribbon(aes(ymin=gam2.5, ymax=gam97.5), alpha=0.2, fill="#4daf4a") +
  geom_ribbon(aes(ymin=gam5, ymax=gam95), alpha=0.3, fill="#4daf4a") +
  geom_ribbon(aes(ymin=gam25, ymax=gam75), alpha=0.4, fill="#4daf4a") +
  geom_line(aes(y=gam50), linewidth=0.8, col="#4daf4a") + geom_point(aes(y=log_idx), pch=1) +
  labs(x="Year", y="GAM log(monarch abundance)") +
  scale_y_continuous(limits=c(16, 20)) +
  scale_x_continuous(breaks=seq(1995, 2020, 5)) +
  geom_vline(xintercept=2011, lty=2, col="gray20")

# all time series
plot_grid(ts_exp, ts_seg, ts_gam, ts_ssm, ncol=2)
ggsave("figure_2.tiff", width = 8.66, height = 6.33, dpi = 600, units = "in")

# get tyc samples
```

```
tyc_samps <- data.frame(exp=exp_out$BUGSoutput$sims.list$tyc,
                        seg=seg_out$BUGSoutput$sims.list$tyc,
                        ssm=ssm_out$BUGSoutput$sims.list$tyc,
                        gam=gam_out$BUGSoutput$sims.list$tyc)

# summarise tyc
exp_30 <- sum(tyc_samps$exp < -30) / 20000
seg_30 <- sum(tyc_samps$seg < -30) / 20000
ssm_30 <- sum(tyc_samps$ssm < -30) / 20000
gam_30 <- sum(tyc_samps$gam < -30) / 20000
exp_50 <- sum(tyc_samps$exp < -50) / 20000
seg_50 <- sum(tyc_samps$seg < -50) / 20000
ssm_50 <- sum(tyc_samps$ssm < -50) / 20000
gam_50 <- sum(tyc_samps$gam < -50) / 20000
exp_med <- median(tyc_samps$exp)
seg_med <- median(tyc_samps$seg)
ssm_med <- median(tyc_samps$ssm)
gam_med <- median(tyc_samps$gam)

# model summaries
d_sum <- data.frame(model=rep(c("EXP", "SEG", "GAM", "SSM"), 5),
                    metric=c(rep("Deviance", 4),
                             rep("Correlation", 4),
                             rep("LOOIC", 4),
                             rep("Probability -50%", 4),
                             rep("Median TYC", 4)),
                    value=c(dev_exp, dev_seg, dev_gam, dev_ssm,
                            cor_exp, cor_seg, cor_gam, cor_ssm,
                            loic_exp, loic_seg, loic_gam, loic_ssm,
                            exp_50, seg_50, gam_50, ssm_50,
                            exp_med, seg_med, gam_med, ssm_med)) %>%
  mutate(model=factor(model, levels=c("EXP", "SEG", "GAM", "SSM")))

# inset
sum_iset <- d_sum %>% filter(metric=="Median TYC" |
                               metric=="Probability -50%" |
                               metric=="LOOIC") %>%
  ggplot() +
  geom_point(aes(x=model, y=value, color=model), size=3) +
  facet_wrap(~metric, scales="free", nrow=1) +
  labs(y="", x="") + theme(strip.background = element_rect(fill=NA)) +
  scale_fill_manual("Model", values=c("#e41a1c","#984ea3","#4daf4a","#377eb8"))

# tyc histograms
h_all <- ggplot() +
  geom_density(data=tyc_samps, aes(x=exp),
               alpha=0.6, fill="#e41a1c", col="#e41a1c", linewidth=NA) +
  geom_density(data=tyc_samps, aes(x=seg),
               alpha=0.6, fill="#984ea3", col="#984ea3", linewidth=NA) +
  geom_density(data=tyc_samps, aes(x=ssm), alpha=0.6,
               fill="#4daf4a", col="#4daf4a", linewidth=NA) +
  geom_density(data=tyc_samps, aes(x=gam),
               alpha=0.6, fill="#377eb8", col="#377eb8", linewidth=NA) +
  labs(x="Ten-year change estimate (%)", y="Density") +
  scale_x_continuous(limits=c(-100, 300), breaks=seq(-100, 300, 50)) +
  geom_vline(xintercept=-50, lty=2, col="gray20")

# plot tyc posteriors
h_inset <- ggdraw(h_all) + draw_plot(sum_iset, .26, .39, .72, .55)
ggsave("figure_3.tiff", width = 8.66, height = 4.33, dpi = 600, units = "in")
```

```
# model weights
stack_wts <- loo_model_weights(x=list(exp_mod=exp_ll, seg_mod=seg_ll,
                                      ssm_mod=ssm_ll, gam_mod=gam_ll),
                               r_eff_list=list(exp_reff, seg_reff, ssm_reff,
                                               gam_reff))
bma_wts <- loo_model_weights(x=list(exp_mod=exp_ll, seg_mod=seg_ll,
                                    ssm_mod=ssm_ll, gam_mod=gam_ll),
                             r_eff_list=list(exp_reff, seg_reff,
                                             ssm_reff, gam_reff),
                             method = "pseudobma")
mean_wts <- (stack_wts + bma_wts) / 2

# create an aggregate distribution based on weights
get_weighted_posterior <- function(dists, wts, summa=T, histo=T){
  nposts <- length(dists)
  nsamps <- length(dists[[1]])
  nwts <- length(wts)
  samp_mat <- matrix(unlist(dists), nrow=nsamps, ncol=nposts)
  dist_wts <- unlist(wts)
  new_dist <- numeric()
  for(i in 1:nposts){
    subsamp <- sample(samp_mat[,i], size=(nsamps * dist_wts[i]))
    new_dist <- c(new_dist, subsamp)
  }
  summ <- NULL
  if(summa==T){
    summ <- c(mean=mean(new_dist), sd=sd(new_dist),
              quantile(new_dist, probs=c(0.025, 0.05, 0.1, 0.25, 0.5, 0.75, 0.9,
                                         0.95, 0.975)))
  }
  if(histo==T) hist(new_dist)
  return(list(weighted_distribution=new_dist, weights=dist_wts, summaries=summ))
}

mod_wtd_tyc_post <- get_weighted_posterior(dists=as.list(tyc_samps),
                                           wts=as.numeric(mean_wts),
                                           summa=T, histo=F)
mod_wtd_tyc_post$summaries
quantile(mod_wtd_tyc_post$weighted_distribution, probs=seq(0.01, 0.99, 0.01))
mod_wtd_tyc_post$weighted_distribution
sum(mod_wtd_tyc_post$weighted_distribution < -30) /
  length(mod_wtd_tyc_post$weighted_distribution)
sum(mod_wtd_tyc_post$weighted_distribution < -50) /
  length(mod_wtd_tyc_post$weighted_distribution)
# ------------------------------------------------------------------------------


# forecast ssm -----------------------------------------------------------------
# write jags model
cat(file = "ssm_mod.txt", "
model {
### priors
  mu[1] ~ dnorm(y[1], ((0.67-0.21)/1.97)^-2)
  b0 ~ dnorm(0, 0.001)
  b1 ~ dnorm(0, 0.001)
  tau.obs <- 1 / (sigma.obs * sigma.obs)
  sigma.obs ~ dnorm(0.44, ((0.67-0.21)/1.97)^-2)
  tau.proc <- 1 / (sigma.proc * sigma.proc)
  sigma.proc ~ dunif(0, 100)
### likelihood
  for (i in 2:n){
```

```
    y[i] ~ dnorm(mu[i], tau.obs)
    mu[i] <- mu[i - 1] + rate[i]
    rate[i] ~ dnorm(b0 + b1 * (mu[i - 1]), tau.proc)
  }
### derived
  tyc <- ((exp(mu[29]) - exp(mu[19])) / exp(mu[19])) * 100
  for(i in 4:n){
    loglik[i] <- logdensity.norm(y[i], mu[i], tau.obs)
  }
}
")

# bundle jags data
add1 <- 20
y2 <- c(md1$log_idx, rep(NA, add1))
n_years2 <- length(y2)
dat_ssm2 <- list(y = y2, n = n_years2)

# parameters to monitor
parameters <- c("mu")

# call jags and summarise posteriors
ssm_out2 <- jags(dat_ssm2, inits=NULL, parameters, "ssm_mod.txt",
                 n.chains=nc, n.thin=nt, n.iter=ni, n.burnin=nb)

# check a few diagnostics
(ssm_sum2 <- MCMCsummary(ssm_out2))
MCMCtrace(ssm_out2, "mu[5]", ISB=F, pdf=F)
MCMCtrace(ssm_out2, "mu[10]", ISB=F, pdf=F)
MCMCtrace(ssm_out2, "mu[15]", ISB=F,pdf=F)
MCMCtrace(ssm_out2, "mu[49]", ISB=F,pdf=F)

# get draws for mu[49]
samps_49 <- as.numeric(MCMCchains(ssm_out2, "mu[49]", ISB = F))

# summarise mu49
summary(samps_49)
p_200k <- sum(samps_49 < log(200000)) / length(samps_49)
p_1m <- sum(samps_49 < log(1000000)) / length(samps_49)
p_3m <- sum(samps_49 < log(3000000)) / length(samps_49)
p_5m <- sum(samps_49 < log(5000000)) / length(samps_49)
p_200k; p_1m; p_3m; p_5m
sum(samps_49 < log(12800000)) / length(samps_49)

# mu 49 histogram
hist_49 <- ggplot() +
  geom_density(data=data.frame(x=samps_49), aes(x=x),
               alpha=0.6, fill="#984ea3", col="#984ea3", linewidth=NA) +
  labs(x="Estimated log(monarch abundance) in 2041", y="Density") +
  scale_x_continuous(limits=c(12, 23), breaks=seq(5, 30, 1)) +
  geom_vline(xintercept=log(200000), lty=5, col="gray10") +
  geom_vline(xintercept=log(1000000), lty=5, col="gray30") +
  geom_vline(xintercept=log(3000000), lty=5, col="gray50") +
  geom_vline(xintercept=log(5000000), lty=5, col="gray70")
hist_49
ggsave("figure_4.tiff", width = 8.66, height = 4.33, dpi = 600, units = "in")

# compute fitted values
ssm_mus2 <- as.data.frame(t(apply(t(ssm_out2$BUGSoutput$sims.list$mu), 1, quantile,
                                  probs=c(0.025, 0.05, 0.25, 0.5, 0.75, 0.95, 0.975)))) %>%
  rename_with(~ paste0("ssm", gsub("%", "", .)))
```

```
md2 <- md1 %>% select(1,4) %>% add_row(year=2022:2041, log_idx=NA) %>%
  bind_cols(ssm_mus2)

# make ts plot
ts_ssm2 <- ggplot(md2, aes(x=year))  +
  geom_ribbon(aes(ymin=ssm2.5, ymax=ssm97.5), alpha=0.2, fill="#984ea3") +
  geom_ribbon(aes(ymin=ssm5, ymax=ssm95), alpha=0.3, fill="#984ea3") +
  geom_ribbon(aes(ymin=ssm25, ymax=ssm75), alpha=0.4, fill="#984ea3") +
  geom_line(aes(y=ssm50), linewidth=0.8, col="#984ea3") + geom_point(aes(y=log_idx), pch=1) +
  labs(x="Year", y="SSM log(abundance)") +
  scale_y_continuous(limits=c(12, 20.5)) +
  scale_x_continuous(breaks=seq(1995, 2040, 5)) +
  geom_hline(yintercept=log(200000), lty=2, col="gray10") +
  geom_hline(yintercept=log(1000000), lty=2, col="gray30") +
  geom_hline(yintercept=log(3000000), lty=2, col="gray50") +
  geom_hline(yintercept=log(5000000), lty=2, col="gray60") +
  geom_vline(xintercept=2021, lty=2, col="gray60") +
  geom_vline(xintercept=2041, lty=2, col="gray60")
ts_ssm2
# ------------------------------------------------------------------------------


# density independent ssm -------------------------------------------------------
# write jags model with b1 fixed to 0 for density independence
cat(file = "ssm_mod.txt", "
model {
### priors
  mu[1] ~ dnorm(y[1], ((0.67-0.21)/1.97)^-2)
  b0 ~ dnorm(0, 0.001)
  b1 <- 0 # dnorm(0, 0.001)
  tau.obs <- 1 / (sigma.obs * sigma.obs)
  sigma.obs ~ dnorm(0.44, ((0.67-0.21)/1.97)^-2)
  tau.proc <- 1 / (sigma.proc * sigma.proc)
  sigma.proc ~ dunif(0, 100)
### likelihood
  for (i in 2:n){
    y[i] ~ dnorm(mu[i], tau.obs)
    mu[i] <- mu[i - 1] + rate[i]
    rate[i] ~ dnorm(b0 + b1 * (mu[i - 1]), tau.proc)
  }
### derived
  tyc <- ((exp(mu[29]) - exp(mu[19])) / exp(mu[19])) * 100
  for(i in 4:n){
    loglik[i] <- logdensity.norm(y[i], mu[i], tau.obs)
  }
}
")

# bundle jags data
dat_ssm3 <- list(y = md1$log_idx, n = n_years)

# parameters to monitor
parameters <- c("b0", "rate", "sigma.proc","sigma.obs", "loglik",
                "tyc", "b1", "mu")

# call jags and summarise posteriors
ssm_out3 <- jags(dat_ssm3, inits=NULL, parameters, "ssm_mod.txt",
                 n.chains=nc, n.thin=nt, n.iter=ni, n.burnin=nb)

# check a few diagnostics
(ssm_sum3 <- MCMCsummary(ssm_out3))
```

```
MCMCtrace(ssm_out3, "b0", pdf=F)
MCMCtrace(ssm_out3, "sigma.proc", pdf=F)
MCMCtrace(ssm_out3, "sigma.obs", pdf=F)
MCMCtrace(ssm_out3, "tyc", ISB=T, pdf=F)
MCMCtrace(ssm_out3, "mu[5]", ISB=F, pdf=F)
MCMCtrace(ssm_out3, "mu[10]", ISB=F, pdf=F)
MCMCtrace(ssm_out3, "mu[15]", ISB=F,pdf=F)

# get logliks
ssm_ll3 <- ssm_out3$BUGSoutput$sims.list$loglik
ssm_reff3 <- relative_eff(exp(ssm_ll3), chain_id = rep(1:2, each = 10000))

# compare looic of d-dependent and d-independent ss models
(loic_ssm <- round(loo(ssm_ll, r_eff=ssm_reff)$estimates['looic','Estimate'], 2))
(loic_ssm3 <- round(loo(ssm_ll3, r_eff=ssm_reff3)$estimates['looic','Estimate'], 2))

# confirm density dependence in raw data with standard technique
rt <- diff(md1$log_idx)
Nt <- md1$log_idx[-29]
time <- 1:length(Nt)
plot(rt~Nt); coef(lm(rt~Nt))
# ------------------------------------------------------------------------------
```
